# Genome-wide methylome stability and parental effects in the worldwide distributed Lombardy poplar

**DOI:** 10.1101/2023.03.14.532102

**Authors:** An Vanden Broeck, Tim Meese, Pieter Verschelde, Karen Cox, Berthold Heinze, Dieter Deforce, Ellen De Meester, Filip Van Nieuwerburgh

**Author notes:** These authors contributed equally (An Vanden Broeck, Tim Meese).

## Abstract

**Background:** Despite the increasing number of epigenomic studies in plants, little is known about the forces that shape the methylome in long-lived woody perennials. The Lombardy poplar offers an ideal opportunity to investigate the impact of the individual environmental history of trees on the methylome.

**Results:** We present the results of three interconnected experiments on Lombardy poplar. In the first experiment, we investigated methylome variability during a growing season and across vegetatively reproduced generations. We found that ramets collected over Europe and raised in common conditions have stable methylomes in symmetrical CG-contexts. In contrast, seasonal dynamics occurred in methylation patterns in CHH-context. In the second experiment, we investigated whether methylome patterns of plants grown in a non-parental environment correlate with the parental climate. We did not observe a biological relevant pattern that significantly correlates with the parental climate. Finally, we investigated whether the parental environment has persistent carry-over effects on the vegetative offsprings’ phenotype. We combined new bud set observations of three consecutive growing seasons with former published bud set data. Using a linear mixed effects analysis, we found a statistically significant but weak short-term, parental carry-over effect on the timing of bud set. However, this effect was negligible compared to the direct effects of the offspring environment.

**Conclusions:** Genome-wide cytosine methylation patterns in symmetrical CG-context are stable in Lombardy poplar and appear to be mainly the result of random processes. In this widespread poplar clone, methylation patterns in CG-context can be used as bio-markers to infer a common ancestor and thus to investigate the recent environmental history of a specific Lombardy poplar. The Lombardy poplar shows high phenotypic plasticity in a novel environment which enabled this clonal tree to adapt and survive all over the temperate regions of the world.

## 1 Background

The contribution of epigenetic variation to adaptation and evolution in plants has been a point of debate during the past decades [1, 2]. In contrast to animals, where epigenetic reprogramming occurs in germ cells and early embryos, epigenetic modifications in somatic plant cells are generally stable [3]. As flowers are induced from somatic cells, it has been hypothesized that epigenetic modifications can induce an ecological memory that is maintained over many generations [4]. Among several epigenetic mechanisms, DNA-methylation is the most studied process. It regulates gene expression and silencing of transposable elements. Variation in DNA-methylation can thus affect the plant phenotype by modifying gene expression [5]. DNA-methylation patterns can be stably transmitted over two or more plant generations [6]. However, it is unclear to which degree inherited epigenetic variation is randomly induced or due to environmental conditions experienced during the parental generation [7]. Particularly for long-lived woody perennials, information on inherited genome-wide methylation patterns is scarce [8]. For example, little is known on how genome-wide DNA methylation patterns (randomly induced or not) vary in woody perennials sharing the same habitat and/or population history. Also, there is a need for better understanding the impact of transmitted epigenetic patterns on the plant phenotype. Neither the magnitude nor the mechanisms behind parental effects are fully understood.

Inherited epigenetic patterns have also been proposed as an explanation for the “missing heritability” [9]. Genome-wide association studies (GWAS) of complex plant traits, such as bud set in trees, consistently found that genetic factors only explained a minority of the expected heritable fraction [10]. This unidentified heritable fraction is called the “missing heritability” [9]. Theoretical frameworks have been developed that quantify the effect of “missing heritability” on complex traits [5, 11] but little empirical information currently exists [1, 5]. For empirically investigating the fitness consequences of epigenetic inheritance, it is important to clearly distinguish between genetic, epigenetic and other sources of transgenerational variation [1]. Genetic replicates (i.e. clonal copies) provide ideal systems to investigate the potential role of epigenetic inheritance in adaptive evolution [1].

The Lombardy poplar (*Populus nigra* cv. ‘Italica’ Duroi) is an excellent study system to investigate how long-lived woody perennials with a prevailing vegetative reproduction can cope with widely contrasting environmental conditions, without variation at the genetic level [12]. This clonal variety of *P. nigra* L. originates from a single mutant male tree and has been distributed worldwide by humans since the beginning of the 18th century [13–15]. The Lombardy poplar is propagated by cuttings from plant material that has been grown locally, sometimes for centuries. It can thus be expected that the large-scale vegetative reproduction of this cultivar in space and time may have resulted in the accumulation of lineage-specific epimutations. Further, some of these epimutations might be non-neutral and environmentally-driven. Such environmentally-driven epigenetic changes could potentially affect the plant phenotype and, when transmitted to vegetative offspring, result in transgenerational phenotypic plasticity. Indeed, in the one-year-old vegetative offspring of the clonal Lombardy poplar and thus in absence of genetic variation, a former study found significant variation in the timing of bud set in a common environment experiment [12]. Ramets (i.e. vegetative offspring of a single parent or ortet) collected from Lombardy poplar ortets from colder origins set bud slightly quicker than ramets obtained from ortets from warmer origins [12]. Whether this transgenerational phenotypic effect remains stable over several years is not known. Based on a relatively small number of methylation-sensitive amplified fragment length polymorphisms (MS-AFLP), the latter study could not directly link epigenetic variation with the climate at the home site of the parent-of-origin or with the observed variation in bud set phenology [12]. In different (non-clonal) study species however, and using more powerful whole-genome bisulfite-sequencing (WGBS) methodologies, associations were found at the single-nucleotide methylation level with spatial structure and climate variables in the long-lived *Quercus lobata* [16, 17], in the annual herb *Arabidopsis thaliana* [18–20], in *Fragaria vesca* [21] and even in the Lombardy poplar [22]. Such observational studies associate epigenetic variation with the environment but they cannot determine whether epigenetic variation is environmentally induced or spontaneous. Most former plant studies based on genome-wide scans and focused on detecting differentially methylated regions (DMRs) among individuals use predefined regions or fixed sliding windows to identify DMRs [e.g. 22]. This latter approach is not suitable for identifying DMRs of arbitrary size by WGBS and does not control for false positives adequately [23]. The sliding windows-approach may result in loss of power or in misleading conclusions on potential functional roles of DMRs in the regulation of gene expression [23]. Another limitation of previous plant methylome studies has been that they did not assess uncertainty and ignore correlation across loci or biological variability from sample to sample [23].

Here, we present the results of three interconnected studies on Lombardy poplar; two observational experiments on the methylome and one study on phenotypic plasticity in a common environment experiment. In the first methylome experiment, we investigated methylome dynamics during early plant life. We studied the variability of the methylome during a growing season and across vegetatively reproduced generations. In the second methylome experiment, we investigated whether methylome patterns of plants grown in a non-parental environment correlate with the parental climate. We searched for data-driven de novo DMRs and DMRs in predefined regions among plants grouped per parental climate. Finally, in the third experiment, we investigated whether the parental environment has persistent carry-over effects on the vegetative offspring’s’ phenotype. We observed timing of bud set in a common environment. We combined new bud set observations of three consecutive growing seasons with former published bud set data [12]. Using a linear mixed effects analysis on the total bud set data spanning four years we investigated whether the observed transgenerational phenotypically plastic effect in terms of timing of bud set persisted.

We found considerable variation between the methylomes of Lombardy poplar ramets collected on ortets growing over Europe and grown in a common environment. DNA methylation in symmetrical CG-context was stable across the growing season and stably inherited by vegetative offspring. Remarkably, these stable, likely neutral methylation patterns can be used as a biomarker for tracing a recent common ancestor of genetically identical ramets. In contrast, seasonal methylation dynamics occurred in CHH-context. We observed a weak, short-term parental carry-over effect on the timing of bud set. However, we did not detect correlations between variation in DNA methylation and the parental climate. This study provides new insights in parental effects and in methylome stability within and across growing seasons and across asexual generations in a long-lived, woody perennial. The results contribute to the understanding of transgenerational plasticity and provide further insight into the biological significance of epigenetic diversity in plants.

## 2 Results

### 2.1 Experimental set-up

The Lombardy poplar represents one single, male genotype that can be recognized by its narrow, columnar growth form (Figure 1A). However, since the beginning of the 20^th^ century, the Lombardy poplar was used in poplar breeding programs with controlled crosses resulting in several other fastigiate cultivars of *Populus nigra* e.g. *P. nigra* cv. ‘Thevestina’ Dode [15]. We therefore selected plant material for this study that was described and verified as the ‘true’ Lombardy poplar previously [12, 24] thus representing one single genotype.

**Figure 1.**
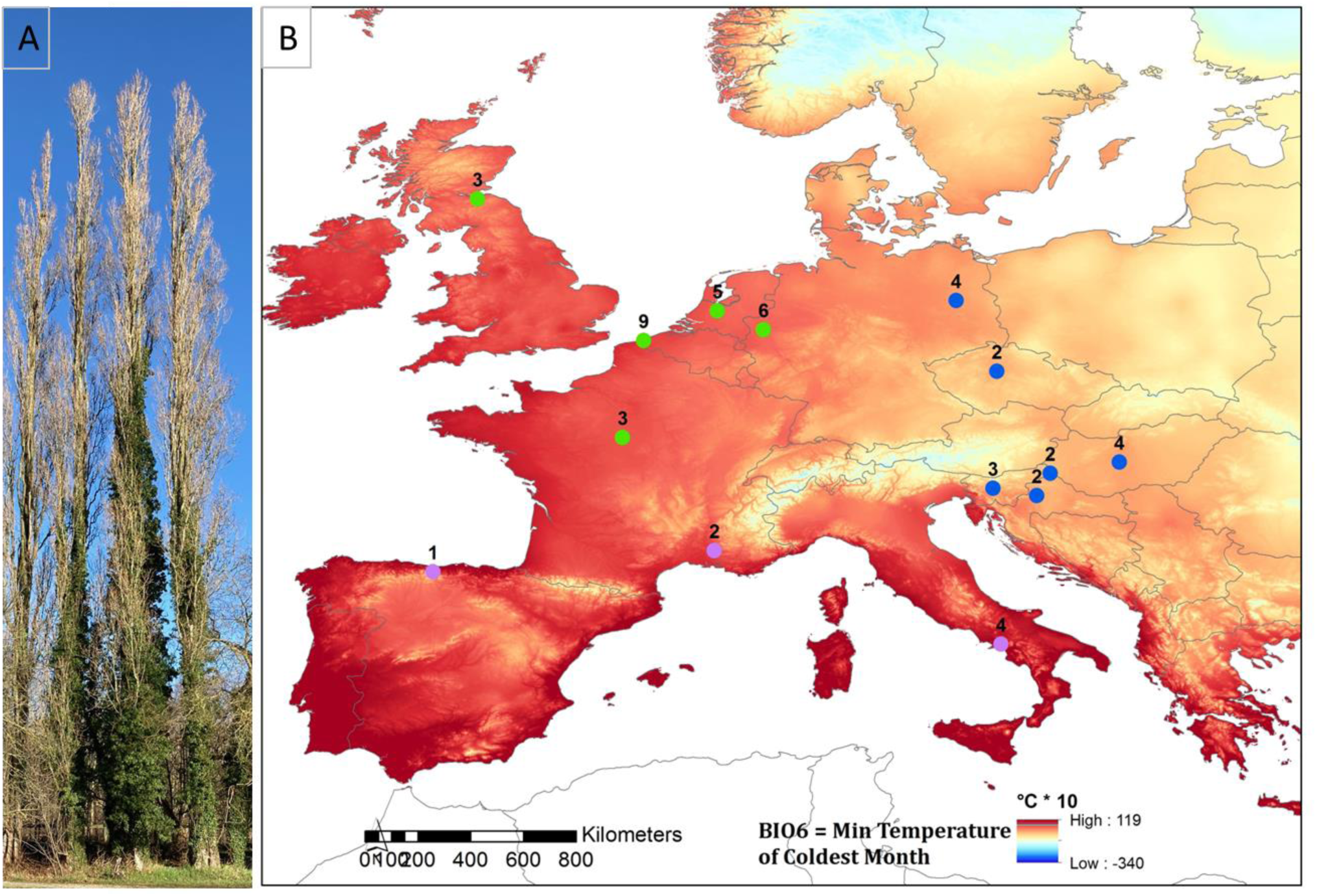
**A. Lombardy poplars. B. Location of the collection sites of the Lombardy poplar ramets included in the DNA pools.** The number of trees included in each DNA pool is indicated on the map. The DNA pools were categorized in three groups for the analysis of differential methylated regions. The grouping was based on the mean temperature in January at the location of the Lombardy poplar ortets calculated for the period 1965 – 2015 (data from CRU TS Version 4.05). (green; 1.83°C – 3.41°C; blue; −1.40°C – −1.18°C, purple; 4.97°C – 8.02°C).

We performed three studies on Lombardy poplar; two observational studies on the methylome and one phenotyping study in a common environment experiment. In the first methylome study (Figure 2A), we investigated methylome dynamics during early plant life. We investigated methylomes of 16 individual plants (Table 1) using WGBS. The plants were grown in a common greenhouse environment in Geraardsbergen, Belgium (lat. 50,77635°, lon. 3.881007°). We used DNA samples of individual plants to investigate the stability of the epigenome during a growing season and across vegetatively reproduced generations. The DNA samples from individual plants included i) DNA-samples from first generation vegetative offspring (F_1_-ramets) collected from Lombardy poplars located in Hungary, Italy, Spain and the UK, and ii) DNA-samples from second-generation vegetative offspring (F_2_-ramets), propagated from the former F_1_-ramets. We extracted DNA from one single leaf of the F_1_-ramets collected in 2017 at the beginning (March 31) and near the end (July 25) of the growing season. Vegetative propagation of four F_1_-ramets resulted in one to three F_2_-ramets per propagated F_1_-ramet. Of these, we collected one leaf on June 12, 2018 for DNA-extraction and WGBS. In the second methylome study (Figure 2B) we investigated whether methylome variation within plants grown in a non-parental environment correlates with the parental climate. We performed WGBS on 13 DNA pools. We used DNA pools to investigate whether epigenetic variation within vegetative offspring in a non-parental environment correlates with parental climatic conditions. We prepared 13 pools of DNA by mixing the same amount of DNA from 2 to 9 (mean: 3.6) individuals. The pools were composed of F_1_-ramets grown in a common environment in Geraardsbergen (Belgium) and originally collected on Lombardy poplar ortets growing in proximity (< 40 km). These F_1_-ramets were collected on in total 53 Lombardy poplar ortets growing across Europe. We used the average January temperature at the geographic location of the ortet as a proxy for the parental climate. We sequenced pools to reduce inter-individual methylation variation and fluctuations in methylation rate caused by stochastic coverage or read sampling bias [2]. The locations of the sampled adult Lombardy poplars (i.e. ortets) are presented in Figure 1B. Additional file 1 includes detailed information on the Lombardy poplars sampled for WGBS.

**Figure 2.**
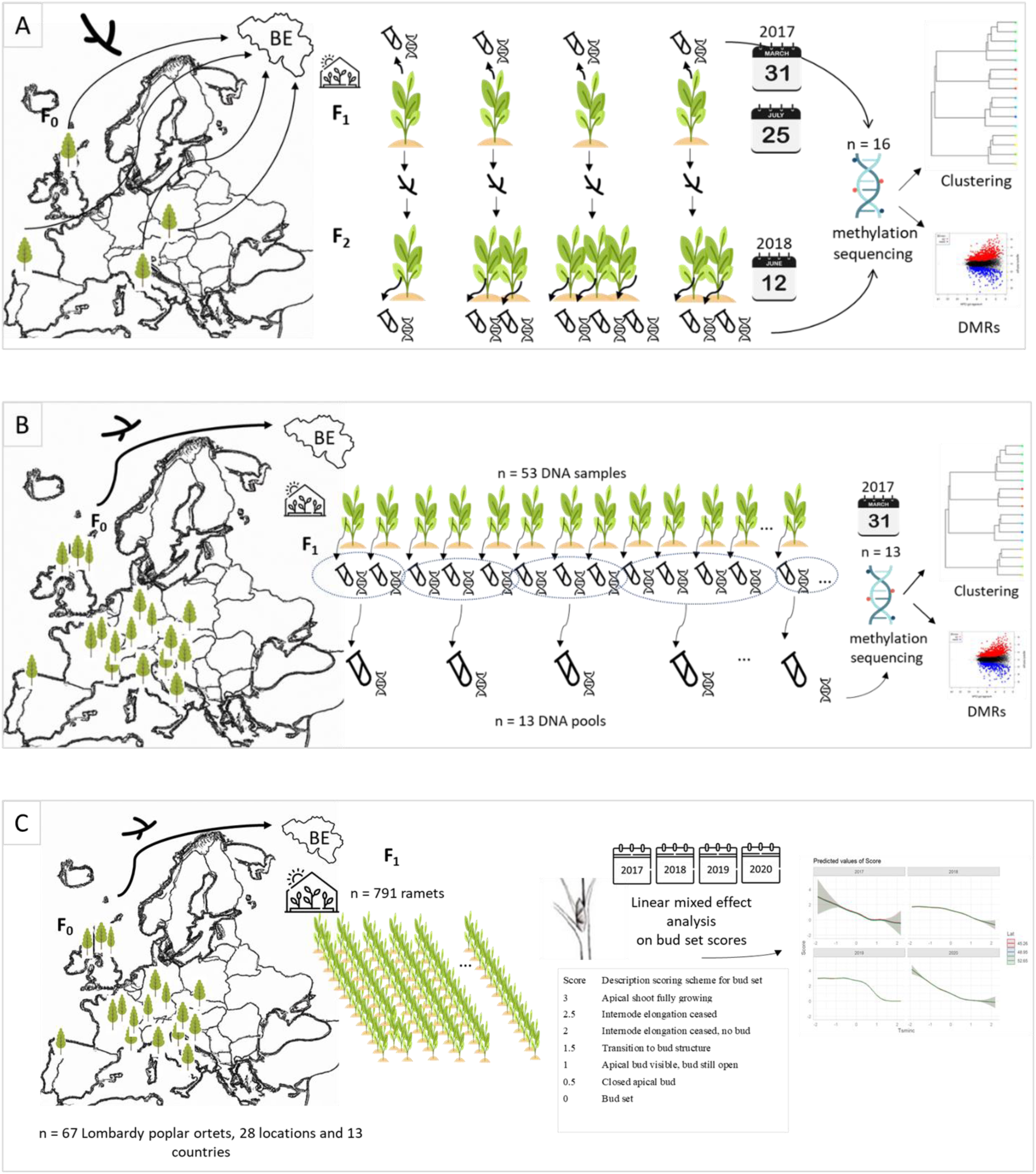
**A. Experimental set-up of the first methylome study.** We sampled 16 plants of Lombardy poplar grown in a common environment in Belgium. These plants included first-generation vegetative offspring (F_1_) grown from ramets collected from four adult Lombardy poplars (F_0_) located in Hungary, Italy, Spain and the UK, respectively, and second-generation vegetative offspring (F_2_). We took leaf samples in 2017 on March 31 and July 25, and in 2018 on June 12. We extracted DNA from one leaf per plant and performed whole genome bisulfite sequencing (WGBS). We used clustering analysis to study the methylome stability across a growing season and across vegetative generations. We studied differentially methylated regions (DMRs) by grouping the WGBS data by the F_0_-parent. **B. Experimental set-up of the second methylome study.** We sampled 53 plants of Lombardy poplar grown in a common environment in Belgium. These plants included first-generation vegetative offspring (F_1_) grown from ramets collected from 53 adult Lombardy poplars (F_0_) located over Europe. We took leaf samples in March 2017 and we extracted DNA from one leaf per plant. We prepared 13 DNA-pools from 2 to 9 F_1_-plants originating from F_0_ -Lombardy poplars growing in proximity. We performed WGBS and searched for DMRs by grouping the WGBS data based on the climate of the F_0_-parent, using the average January temperature as a proxy for the parental climate, resulting in three groups. **C. Experimental set-up of the common environment experiment.** We observed timing of bud set on 791 plants of Lombardy poplar grown in a common environment in Belgium using a seven-stage scoring. These plants included F_1_-ramets collected from 67 adult Lombardy poplars (F_0_) grown on 28 different locations in 13 different countries over Europe. We used a linear mixed effects analysis on the total bud set data spanning four years (2017 – 2020) to investigate the relationship between bud set phenology of the ramets in the common environment experiment (F_1_) and the latitude of origin of the home site of the Lombardy poplar parent-of-origin (F_0_). Icons made by Freepik from www.flaticon.com.

**Table 1.** Leaf samples collected for the first methylome study from individual Lombardy poplar ramets.

| ID_Leaf Sample | ID_WGBS | Ramet | Offspring | Sampling |  | Ortet's |  |  |  |
| --- | --- | --- | --- | --- | --- | --- | --- | --- | --- |
|  |  |  |  | Date | Ortet | Location | Latitude | Longitude | Elevation (m) |
| HUN4_F1_March2017 | WGBS01 | HUN4014 | F<sub>1</sub> | 31/03/2017 | HUN4 | Hungary | 46,93713 | 18,80150 | 137 |
| HUN4_F1_July2017 | WGBS05 | HUN4014 | F<sub>1</sub> | 25/07/2017 | HUN4 | Hungary | 46,93713 | 18,80150 | 137 |
| HUN4_F2_June2018 | WGBS14 | HUN4014_01 | F<sub>2</sub> | 12/06/2018 | HUN4 | Hungary | 46,93713 | 18,80150 | 137 |
| ITS3_F1_March2017 | WGBS02 | ITS3014 | F<sub>1</sub> | 31/03/2017 | ITS3 | Italy | 40,75182 | 14,77437 | 207 |
| ITS3_F1_July2017 | WGBS06 | ITS3014 | F<sub>1</sub> | 25/07/2017 | ITS3 | Italy | 40,75182 | 14,77437 | 207 |
| ITS3_F2_01_June2018 | WGBS10 | ITS3014_01 | F<sub>2</sub> | 12/06/2018 | ITS3 | Italy | 40,75182 | 14,77437 | 207 |
| ITS3_F2_02_June2018 | WGBS15 | ITS3014_02 | F<sub>2</sub> | 12/06/2018 | ITS3 | Italy | 40,75182 | 14,77437 | 207 |
| SPC1_F1_March2017 | WGBS04 | SPC1001 | F<sub>1</sub> | 31/03/2017 | SPC1 | Spain | 43,21521 | -4,59266 | 279 |
| SPC1_F1_July2017 | WGBS07 | SPC1001 | F<sub>1</sub> | 25/07/2017 | SPC1 | Spain | 43,21521 | -4,59266 | 279 |
| SPC1_F2_02_June2018 | WGBS09 | SPC1001_02 | F<sub>2</sub> | 12/06/2018 | SPC1 | Spain | 43,21521 | -4,59266 | 279 |
| SPC1_F2_01_June2018 | WGBS11 | SPC1001_01 | F<sub>2</sub> | 12/06/2018 | SPC1 | Spain | 43,21521 | -4,59266 | 279 |
| SPC1_F2_03_June2018 | WGBS13 | SPC1001_03 | F<sub>2</sub> | 12/06/2018 | SPC1 | Spain | 43,21521 | -4,59266 | 279 |
| UKD2_F1_March2017 | WGBS03 | UKD2010 | F<sub>1</sub> | 31/03/2017 | UKD2 | UK | 55,89000 | -3,07050 | 57 |
| UKD2_F1_July2017 | WGBS08 | UKD2010 | F<sub>1</sub> | 25/07/2017 | UKD2 | UK | 55,89000 | -3,07050 | 57 |
| UKD2_F2_01_June2018 | WGBS12 | UKD2010_01 | F<sub>2</sub> | 12/06/2018 | UKD2 | UK | 55,89000 | -3,07050 | 57 |
| UKD2_F2_02_June2018 | WGBS16 | UKD2010_02 | F<sub>2</sub> | 12/06/2018 | UKD2 | UK | 55,89000 | -3,07050 | 57 |

Finally, in the third experiment, we investigated whether the parental environment has persistent carry-over effects on the vegetative offspring’s’ phenotype. We observed timing of bud set in a common environment (Figure 2C). We combined new bud set observations of three consecutive growing seasons with former published bud set data [12]. Using a linear mixed effects analysis on the total bud set data spanning four years we investigated whether the parental carry-over effect in terms of timing of bud set persisted.

### 2.2 Epigenetic variation and methylome stability

On average 46% (range 44% - 49%) of the sequence reads obtained by whole genome bisulfite sequencing (WGBS) from the 29 DNA samples were mapped uniquely to the genome of *Populus trichocarpa.* The average percentage of total cytosines that were methylated in the 16 individual Lombardy poplar ramets was 22%, 11% and 2% in the CpG-, CHG- and CHH-context, respectively (where H represents A, C or T). Additional file 2 includes the mapping and methylation statistics for each of the 16 genomes for the three sequence contexts. The percentage of cytosine methylation in the three sequence contexts for each of the 16 individual ramets of Lombardy poplar sampled is given in Figure 3. The dendrogram based on the differentially methylated cytosines in CpG-context as well as in CHG-context from the individual ramets clearly grouped the F_1_- and the F_2_-ramets according to their parent-of-origin (F_0_) (i.e. ortet) (Figure 4A). Similar clustering results were obtained based on the multidimensional scaling analyses (MDA) (Figure 4B). Methylation in symmetrical CG-context was stable within and across seasons. In contrast, the hierarchical cluster analysis clustered the samples based on differentially methylated cytosines in CHH-context according to the sampling period in the growing season; samples taken at the beginning of the growing season (March) clustered clearly separately from the samples taken in the middle of the growing season (June and July). However, the latter pattern was less clear on the scatter plot of the first two dimensions obtained by the MDA.

**Figure 3.**
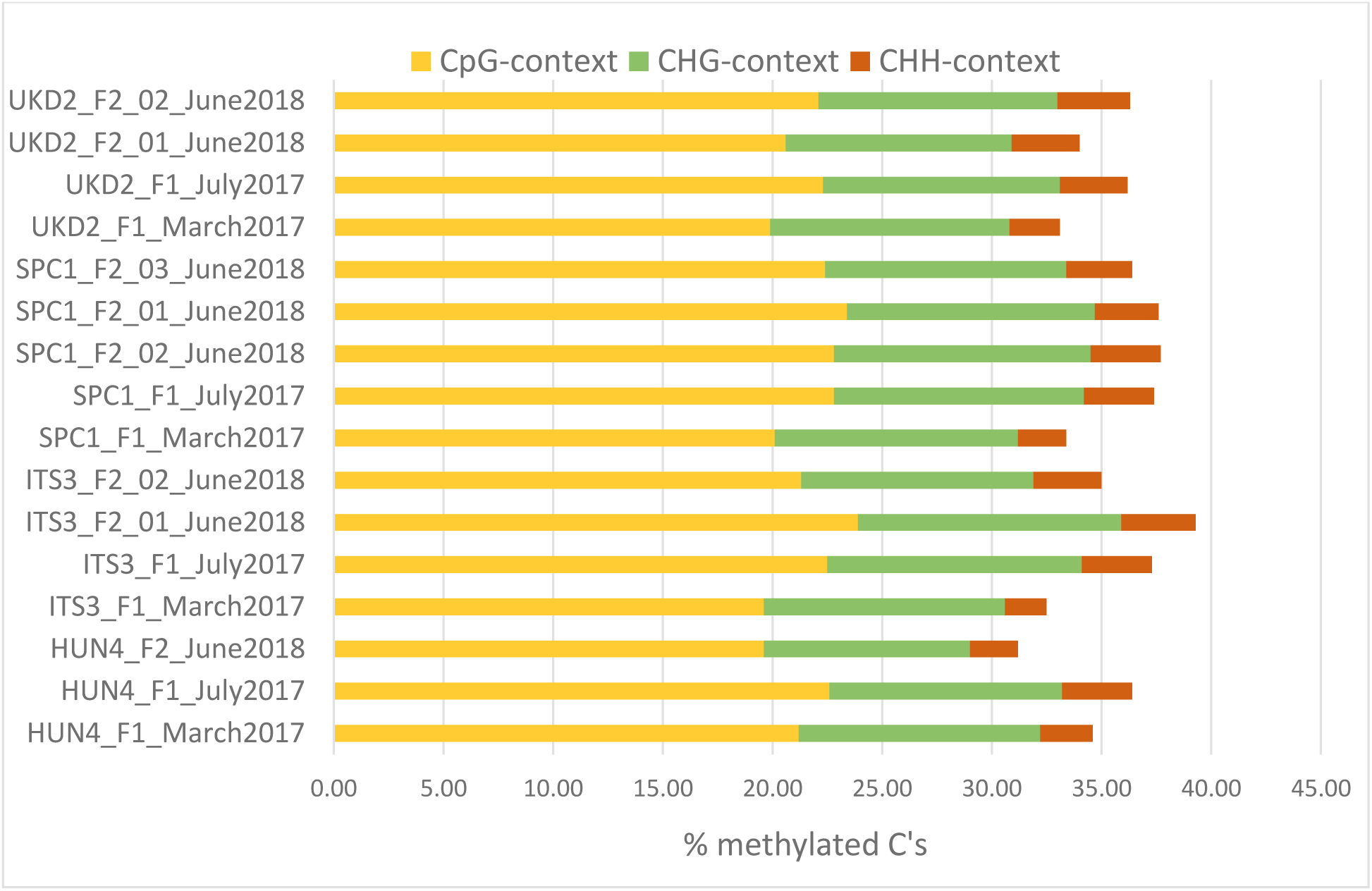
Percentage of cytosine methylation per sequence context for each of the 16 Lombardy poplar ramets. Sample names are explained in Table 1.

**Figure 4.**
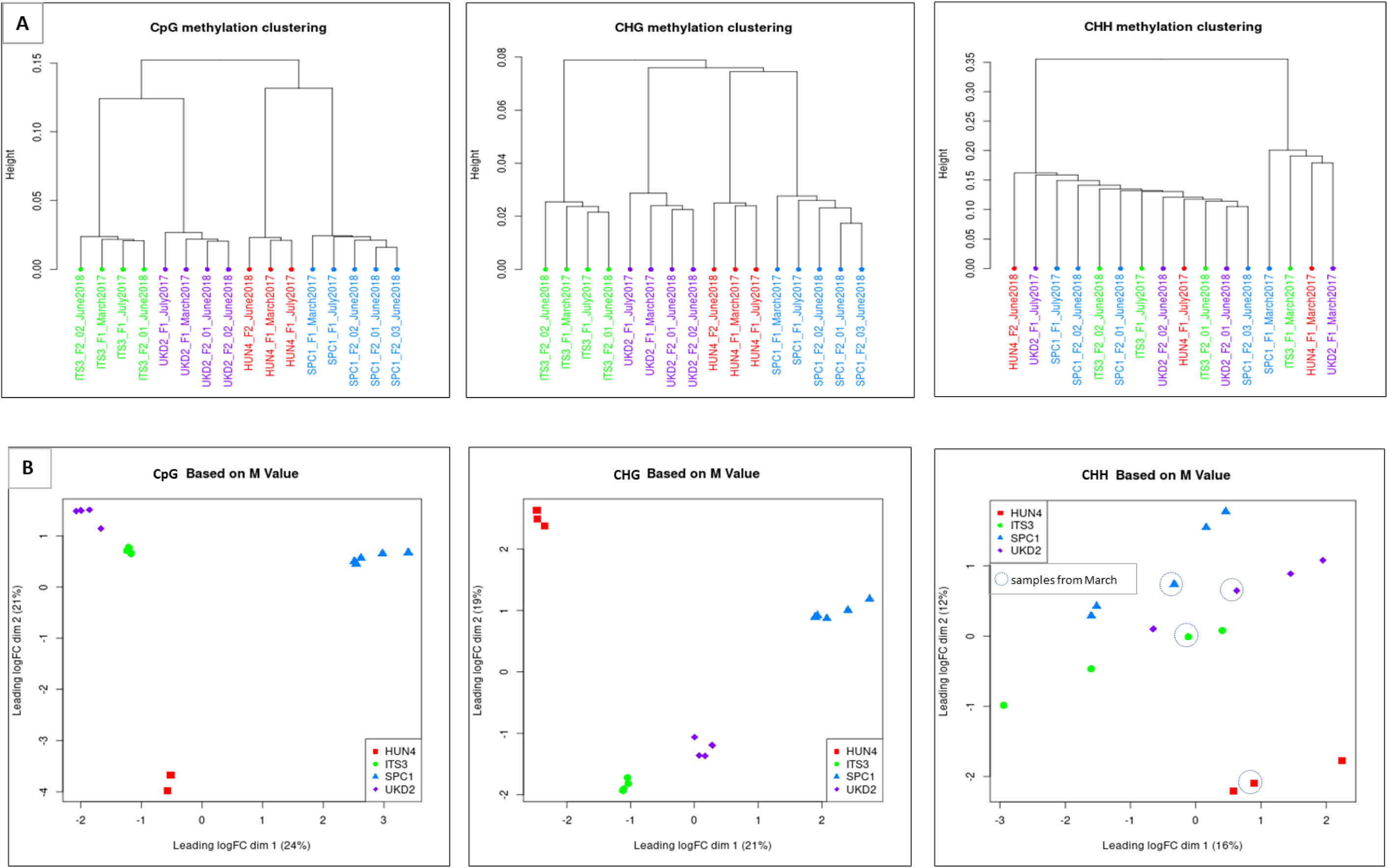
Clustering results based on methylation-states per sequence context for each of the 16 Lombardy poplar ramets. **A. Hierarchical clustering. B. Multidimensional scaling (MDS).** Sample names are explained in Table 1. Different colors represent the four different ortets. Green: ‘ITS3’, purple: ‘UKD2’; red: ‘HUN4’ and blue: ‘SPC1’ located in Italy, UK, Hungary and Spain, respectively.

To explore the functionally more relevant differentially methylated regions (DMRs) compared to DMPs [25], we identified DMRs in predefined regions (gene bodies and promoters). In total, we identified in total 2881 statistically significant DMRs (p-value < 0.05) within the pre-defined genomic gene bodies and promoter regions, for the three sequence contexts (CpG, CHG and CHH-DMRs) between any of the six pairwise comparisons of the individual ramets grouped per ortet. The majority of these DMRs were found in CpG- and CHG-context (53% and 45%, respectively) and within genes (84.4%). Most of the DMRs were common to two or more pairwise comparisons (63% and 74% for promoter and gene regions, respectively) (Figure 5A). A list of all DMRs identified in predefined regions is presented in Additional file 3.

**Figure 5.**
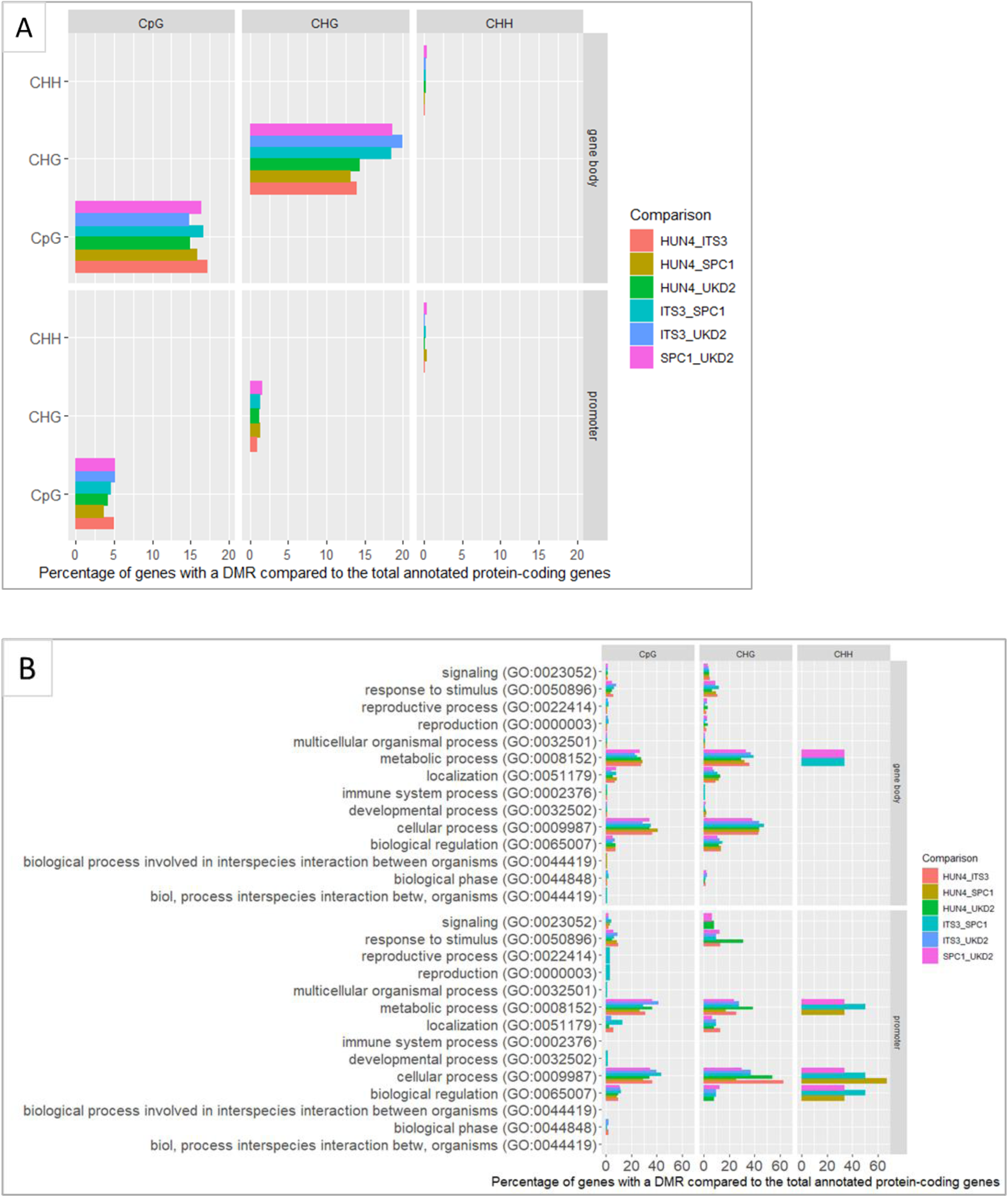
**A. Genes in differentially methylated regions (DMRs) per sequence context in gene bodies and in promoters.** Samples of 16 individual Lombardy poplars grown in a common environment were grouped based on their common ortet; ‘HUN4’, ‘ITS3’, ‘SPC1’, ‘UKD2’ located in Hungary, Italy, Spain and the UK, respectively. Different colors represent different pairwise comparisons between groups. **B. Gene ontology classification**. Bar chart representing the functional categorization of the annotated DMRs (P < 0.05, category; biological process) per sequence context in gene bodies and in promoters. Information on the ortets is given in Table 1.

The overrepresentation of the gene ontology (GO) terms and their biological processes associated with genes located in these DMRs are presented in Figure 5B. GO terms that were enriched in CpG-DMRs located in genes were mainly involved in metabolic and cellular processes (e.g. catalytic activity, receptor binding), whereas the few (0.2%) GO terms in CH-DMRs located in genes were enriched for biological processes involved in regulatory functions (e.g. regulation in gene expression). One gene was differentiated between all pairwise comparisons in CpG-context, a gene with a transferase activity, UBIA PRENYLTRANSFERASE DOMAIN-CONTAINING PROTEIN 1 (PTHR13929:SF0) involved in quinone metabolic process. GO terms that were enriched in CpG-DMRs located in promoters were mainly involved in metabolic and cellular processes (e.g. catalytic activity, receptor binding), whereas few (2%) were involved in biological regulation. Additional file 4 includes the total list of GO terms that were enriched in DMRs. A heatmap of GO terms that were enriched in CpG-DMRs located in promoters per pairwise comparison of ramets grouped per parent-of-origin, is presented in Figure 6. Additional file 5 contains the clustering results and the heatmaps of enriched GO-terms for all pairwise comparisons. No specific DMRs were found associated with rhythmic process (GO:0048511) any process pertinent to the generation and maintenance of rhythms in the physiology of an organism (thus including bud set phenology). There are 17 genes classified within this biological process in *Populus trichocarpa*, none of these occurred in the list of DMRs.

**Figure 6.**
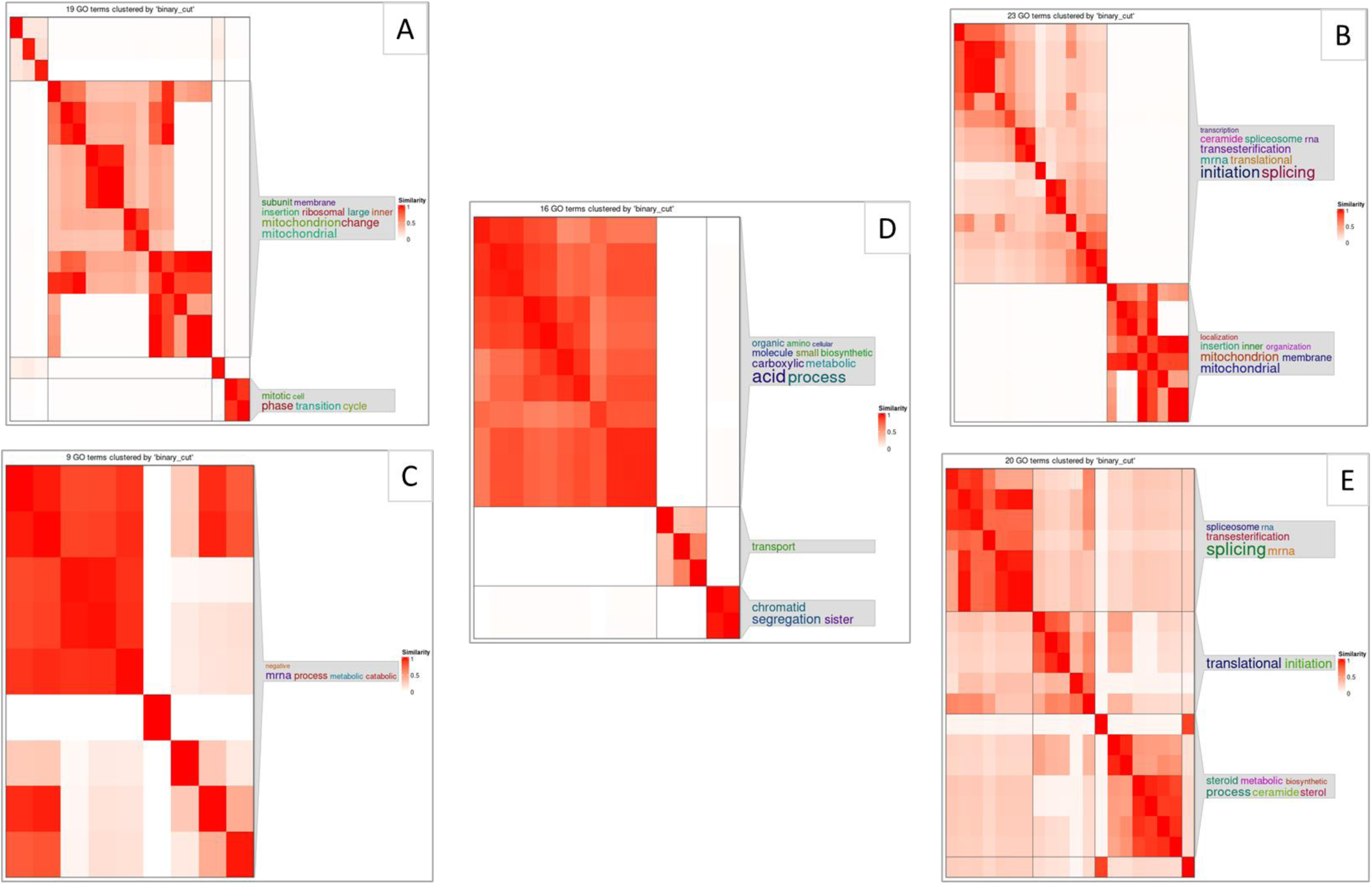
**Heatmaps with GO terms over-represented in promoters containing DMRs in CpG-context per between-group pairwise comparison**. DMRs were identified between groups by grouping the WGBS data from 16 individual Lombardy poplar ramets by their corresponding parent-of-origin (ortet ‘HUN4’ located in Hungary, ‘ITS3’ in Italy, ‘SPC1’ in Spain and ‘UKD2’ in the UK, respectively). **A**. HUN4 versus ITS3; **B.** HUN4 versus UKD2, **C**; ITS3 versus SPC1; **D.** HUN4 versus SCP1, **E**. SPC1 versus UKD2.

In contrast to the WGBS data of the 16 individual ramets, we did not observe any biological relevant pattern in the clustering of the WGBS data of the 13 pooled DNA-samples based on the DMPs. Also, no significant DMRs (neither *de novo,* nor in predefined regions) were detected between pairwise comparisons of the pooled samples grouped based on the average January temperature at the geographic location of the Lombardy poplar parent-of-origin. Moreover, we found no significant *de novo* DMRs when comparing, for each sampling period (March vs. June/July), the methylomes of the individual ramets grouped based on the average January temperature at the geographic location of the Lombardy poplar parent-of-origin (i.e. ramets originated from the ortets from Hungary and UK versus ramets from the ortets located in Spain and Italy).

### 2.3 Variation in timing of bud set

Whether transgenerational phenotypic transmission depends on parent-of-origin is important to understand the mechanisms behind phenotypic plasticity [26]. We grew 791 ramets from 67 adult Lombardy poplar ortets planted across Europe in 28 locations. The ramets were grown in a non-maternal, uniform environment. We analyzed bud set data collected during four consecutive growing seasons. We found a statistical significant but weak short-term, parental carry-over effect of the location of the Lombardy poplar parent-of-origin (in terms of latitude) on the timing of bud set of the ramets in the novel environment. However, this effect was negligible compared to direct effects of the asexual offspring’s environment. There was a statistically significant effect of the latitude-of-origin (Lat) on bud set phenology but the estimated effect was very small and only of biological significance in the first year of the experiment (2017) (Figure 7A). In 2017, moving ramets northward from the latitude of the parent-of-origin, thus into longer days, resulted in a slightly increase in bud set score and thus a slightly longer growth period for these ramets relative to ramets originating from the latitude nearby the experimental site. Although still statistically significant, the effect size of the latitude-of-origin (Lat) is near zero in the years 2018, 2019 and 2020 and of non-biological relevance (Figure 7B). In the first year of the experiment (2017), the effect on bud set phenology of the cumulative daily minimum temperature during plant growth at the experimental site (Tsmin) was about two orders of magnitude larger compared to the effect of the latitude-of-origin (Lat) and thus clearly of higher importance. The effects sizes of the variables affecting the bud set phenology and maintained in the linear mixed effects model are given in Table 2. The response variable was bud set score and the year 2017 was set as the reference year. The values of the estimates given in Table 2 are thus the effect sizes on the response variable compared to the year 2017. The intercept represent the estimate of the response variable for the year 2017, with the values for the variables latitude (Lat) and Tsminc, set to zero. Only two-way interactions were considered. Additional file 6 contains the raw data of the bud set observations and Additional file 7 includes the source codes to reproduce the results of the bud set data analysis.

**Figure 7.**
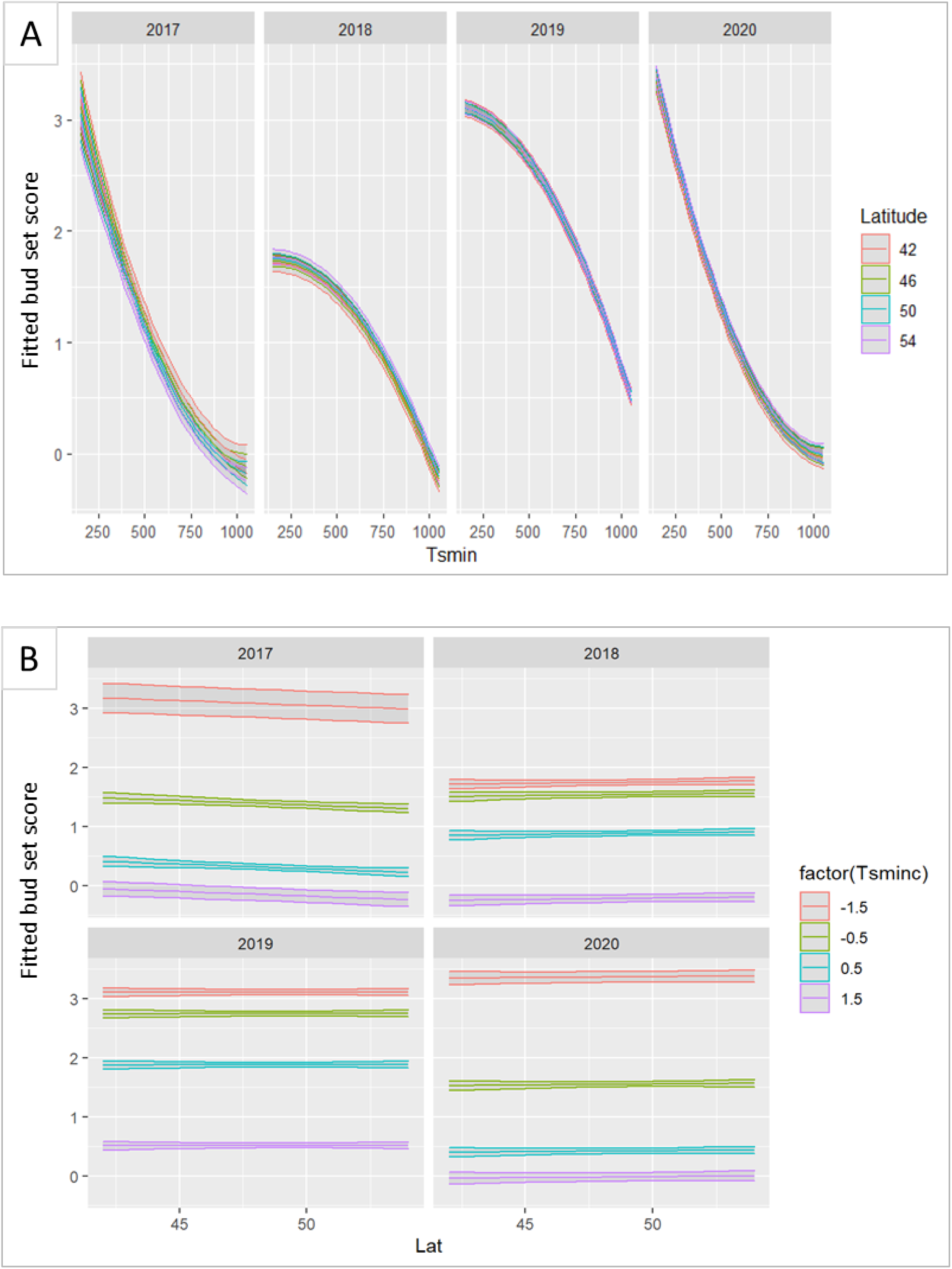
**A. Fitted values for bud set score plotted against the cumulative daily minimum temperature during plant growth.** The ramets originated from Lombardy poplar ortets growing at different latitudes. Bud set score three; apical shoot is fully growing, bud set score zero; the apical bud is formed. Confidence intervals are indicated in grey. **B. Fitted values for bud set score per year plotted against the latitudes of the ortets.** *Tsminc;* the temperature sum of daily minimum temperature at the common environment experiment calculated from 1 July, centered and standardized.

**Table 2.** Effect sizes of the variables maintained in the linear mixed effects model affecting the bud set phenology.

| Parameter | Estimate | Std. Error | z value | p-value |
| --- | --- | --- | --- | --- |
| Intercept | 1.527 | 0.303 | 5.033 | **<0.001** |
| Tsminc | -1.021 | 0.205 | -4.971 | **<0.001** |
| (Tsminc^2) | 0.297 | 0.240 | 1.237 | 0.216 |
| Lat | -0.016 | 0.005 | -3.292 | **0.001** |
| fYear2018 | -0.509 | 0.303 | -1.682 | 0.093 |
| fYear2019 | 0.712 | 0.292 | 2.434 | **0.015** |
| fYear2020 | -0.775 | 0.315 | -2.463 | **0.014** |
| Tsminc:fYear2018 | 0.389 | 0.230 | 1.689 | 0.091 |
| Tsminc:fYear2019 | 0.252 | 0.220 | 1.147 | 0.251 |
| Tsminc:fYear2020 | -0.100 | 0.238 | -0.42 | 0.675 |
| (Tsminc^2):fYear2018 | -0.502 | 0.254 | -1.979 | **0.048** |
| (Tsminc^2):fYear2019 | -0.477 | 0.247 | -1.936 | 0.053 |
| (Tsminc^2):fYear2020 | 0.052 | 0.275 | 0.19 | 0.850 |
| Lat:fYear2018 | 0.020 | 0.003 | 6.612 | **<0.001** |
| Lat:fYear2019 | 0.016 | 0.003 | 5.51 | **<0.001** |
| Lat:fYear2020 | 0.019 | 0.003 | 5.823 | **<0.001** |

*Tsminc*; the temperature sum of daily minimum temperature at the common environment experiment calculated from 1 July, centered and standardized, Lat; the latitude of origin, centered on 50, of the parent- of-origin. Significant p-values (< 0.05) are indicated in bold. p-values of main effects in the presence of an interaction are not to be interpreted but are retained in the model.

## 3 Discussion

### The methylome as a biomarker for origin

The widespread, clonally propagated Lombardy poplar provides an unique opportunity to study epimutation accumulation over long time-scales and in variable environments [12]. This study found several tens of thousands cytosine methylation polymorphisms (DMPs) between genetically identical ramets grown in a common environment and collected on adult trees (ortets) from diverse sites over Europe. Strikingly, ramets grown in the controlled environment for up to two growing seasons continued to retain genome-wide epigenetic marks in symmetrical CG-contexts (i.e. CpG and CHG, where H = A, C, T) typical of the founder tissue used for the initial propagation. The hierarchical cluster analysis grouped individual ramets grown in a novel environment by the parent-of-origin, regardless of the ramets’ sampling date during the growing season or regardless of their asexual reproduction cycle (F_1,_ F_2_). This suggests that genome-wide epigenetic variation in woody perennials in symmetrical CG-contexts is stable and likely mostly random. In contrast, seasonal changes occurred in CHH-methylation. Hierarchical clustering analysis of the individual samples based on DMPs in the asymmetrical CHH-context grouped the samples according to the date of sampling during the growing season (March versus June/July), and thus not by the parent-of-origin. Methylome stability at individual CpG-sites [2] and temperature-dependence variation in methylation at CHH-sites was also found in *Arabidopsis species* [18, 27]. Each of the three contexts of methylation is distinctly regulated and maintained by unique pathways [reviewed by 28, 29]. In plants, methylation occurs mostly in transposable elements and other repetitive DNA-sequences (in CpG, CHG and CHH context) while methylated CG is the most abundant form of cytosine methylation and is often found within transcribed regions (i.e. gene body methylation) [6]. Our findings suggest that in woody plants, geographically structured natural variation in epigenetic patterns in CHH-context may be the result of the variation in seasonality along a latitudinal gradient (i.e. variation in photoperiod and temperature). Thus, these genome-wide methylation patterns in CHH-context reflect phenotypic adjustments to local conditions or intra-individual phenotypic plasticity (i.e. the ability of one genotype to express varying phenotypes when exposed to different environmental conditions [26]) rather than evolutionary adaptation.

The rates of epimutations in poplars are four to five orders of magnitude greater than those of spontaneous somatic mutations. By sampling multiple, age-estimated branches of a single tree of *Populus trichocarpa*, Hofmeister et al. [8] found that symmetrical CG-methylations are cumulative across somatic development and originate mainly from DNA methylation maintenance errors during mitotic cell divisions. In contrast to the strong similarities in per-generation (epi)genetic mutation rates, the per-year (epi)genetic mutation rates are lower in the perennial poplar than in the in the herbaceous, annual plant *Arabidopsis thaliana* [8]. The frequent vegetative reproduction and widespread use of the Lombardy poplar by humans may have facilitated epi-mutation accumulation during mitotic cell divisions. Once established, epimutations can stably persist or be inherited across generations [30]. In a study of white poplar (*Populus alba*), ramets collected on ortets growing at different locations and representing a single clone grouped according to the sampling site based on differentially methylated CG-sites generated by Methylation Sensitive Amplified Fragment Length Polymorphisms (MS-AFLPs) [31]. This is in agreement with the finding of this study that closely related plants sharing a common ancestor show high similarities in genome-wide symmetrical CG-methylation patterns. Epigenetic patterns in symmetrical CG-contexts can thus be used as a bio-marker for identifying a common ancestor or ortet of genetic identical ramets. This may be useful, for example, for providing insights into the range expansion of invasive, clonal plants, like Japanese knotweed. To our knowledge, this is one of the first multigenerational studies that contribute to the recent evidence that spontaneous symmetrical CG-epimutations in long-lived perennials accumulate across asexual generations and at high rate over time [8] resulting in rapid methylome divergence among asexual plant lineages. These CG-epimutations can be used as a fast-ticking molecular clock, not only for age-dating long-lived perennials [8] but also for timing evolutionary events in the recent past [32, 33] like the construction of clonal phylogenies.

Geographically structured epigenetic variation is found in a wide range of organisms [16, 19, 34]. Several observational studies suggest that environment induces epigenetic variation and that selection and local fitness effects have led to the observed epigenetic patterns, [16, 21, 35], but evidence for causal relationships is lacking. Most epimutations are spontaneous and likely neutral [8, 17, 30, 36] though epimutations can result in phenotypic variation in plants [37–39]. In this study no significant differences in methylated regions (DMRs) were detected between pairwise comparisons of the methylomes of pooled samples grouped by average temperature in January at the geographic location of the adult Lombardy poplars. By pairwise comparisons of individual methylomes grouped per parent-of-origin, we found significant DMRs within gene bodies and enriched for cellular and metabolic processes. To date, there is no convincing data to support a direct causal link between the presence/absence of gene body methylation and the modulation of expression in flowering plants [8, 30]. We also found DMRs within promoters that were enriched for biological processes (e.g. biological regulation, response to stimulus) and that could result in transcriptional consequences effecting the plants’ phenotype. In an empirical study testing the effects of drought on transcriptomes in three poplar clones, Raj et al. [40] suggest that clone history can have a profound effect on drought responses. In this study we cannot make conclusions about environmental effects on the epigenome. In trees, local adaptation along climatic gradients is mostly the result of selection on polygenic traits, involving hundreds or thousands of loci, each with a very small effect, that is very difficult to detect [41]. The complex interplay between genotype and environment combined with the limited number of biological replicates, resulted in limited power of this study to identify causal epigenetic variants. However, Zhou et al. [42] detected a functional relationship between DNA-methylation diversity and drought resistance and resilience in *Populus tomentosa* by combining epigenome-wide association studies with genome-wide association studies. Further research is needed building on our findings to help understand the mechanisms by which DNA methylation epialleles are generated, maintained and erased, and their functional consequences in plants.

### Weak, short-term parental effect

Several studies have highlighted the potential role of environmentally induced epigenetic variation on parental effects [43], although empirical evidence is scarce. Here, we experimentally tested the potential effect of epigenetic inheritance on phenotypic variation in terms of bud set in a common garden experiment. We found a significant but very weak and short-term parental carry-over effect on the timing of bud set in the novel environment of vegetative offspring for the Lombardy poplar. The direction of this effect correlated with the latitude-of-origin of the Lombardy poplars; ramets of Lombardy poplars from more southern locations set bud slightly later in summer than trees from more northern locations [12]. Indeed, the movement of a locally adapted poplar genotype from the latitude of origin towards more northern locations, thus into longer days during the growing season, prolongs the period of its active growth while the contrary is observed for movements towards more southern locations [44, 45]. However, the temperature at the new environment had a much stronger influence on the duration of bud formation compared to the parental effect. Furthermore, the weak parental effect on bud set was reduced to near-zero in the three consecutive growing seasons and had little biological relevance after the first growing season. This indicates that the parental effect is very limited compared with the direct effect of the offspring environment and that it becomes negligible after the first growing season. It may therefore rather be an intergenerational effect than a truly transgenerational effect [46]. This short-time epigenetic memory observed in this study is therefore unlikely to be responsible for a contribution to heritable variation, the so-called ‘missing heritability’ or the contribution to phenotypic variation in bud set observed in experimental studies that could not be attributed to the genetic architecture [47, 48]. The results of this study are in agreement with the conclusions of Uller et al. [49] based on a meta-analysis of experimental studies on anticipatory parental effects in plant and animals indicating that parental effects are generally small compared to direct effects of offspring environment, and more subtle than typically assumed. The results of this study show a very high phenotypic plasticity of Lombardy poplar. A rapid response mechanism in bud phenology likely enabled this clonal tree to adapt and survive after human introduction all over the temperate regions of the world and even in subtropical environments.

## 4 Conclusions

The clone history of Lombardy poplar shapes the genome-wide methylome which can be used as biomarkers to identify a common ancestor of genetically identical trees on short evolutionary timescales. Our results show a rapid plastic response of the Lombardy poplar and a weak short-time memory effect on the timing of bud set of the parental environmental cues.

## 5 Methods

### 5.1 Whole Genome Bisulfite Sequencing

#### 5.1.1 DNA extraction

Total genomic DNA was extracted from fresh leaves with the Qiagen Plant DNA kit (Hilden, Germany). The integrity of the DNA was assessed on 1.5% agarose gels, and DNA quantification was performed with Quant-iT™ PicoGreen® dsDNA Assay Kit (Thermo Fisher Scientific, Massachusetts, USA) using a Synergy HT plate reader (BioTek, Vermont, USA).

#### 5.1.2 Library construction and whole genome bisulfite sequencing

Up to 250 ng of DNA was fragmented to 300 bp with a Covaris S2 sonicator prior to whole-genome library preparation (NEBNExt Ultra II kit), ligation of methylated adaptors and size selection on 2% E-gels (400-550 bp). Bisulfite conversion was performed using the EZ DNA methylation gold kit (Zymo Research), followed by an enrichment PCR of 12 cycli. Paired-end 100-bp sequencing of the library fragments was performed on an Illumina Novaseq S2 sequencer, generating on average 83,886,466 ± 14,181,156 reads (mean ± SD) and an average read depth of approximately 17x. The raw fastq datafiles, the processed data and the metadata are available at the [Gene Expression Omnibus (GEO) database](https://www.ncbi.nlm.nih.gov/geo/) (submission GSE225596).

#### 5.1.3 Identification of methylated sites

The raw fastq-files were first quality-based trimmed and adapter trimmed with Trim Galore v0.6.6 (https://doi.org/10.5281/zenodo.5127899). The trimmed reads were subsequently mapped onto the genome of *Populus trichocarpa* (Pop_tri_v3, Ensembl release 48) using Bismark v0.23.0 [50]. As this is the genome of a related species, the mapping parameters were relaxed by specifying the function --score_min L,0,-0.4. Afterwards, the aligned BAM files were deduplicated using Bismark. Lastly, bismark_methylation_extractor and bismark2bedGraph v0.24.0 were used to generate the context-specific methylation-states (i.e. CpG, CHG and CHH) at the individual cytosine-level for each sample. Only loci that were covered in all samples were retained for further analysis. Additional file 8 represents the different steps of the bioinformatics pipeline (i.e. snakefile). Because it is unclear whether differentially methylated positions have any functional consequences in plants [25], we identified methylated regions.

#### 5.1.4 Identification of differential methylated predefined regions

Plant DNA methylation occurs through the link of a methyl group (-CH3) at cytosine in symmetric context; CpG and CHG, and asymmetric sequence context; CHH (where H is any nucleotide except G). We first summarized the identified methylated sites into known regions; promotors and gene bodies. Thereafter, we tested these predefined regions for differential methylation. The value of this approach has been demonstrated before (see [2]). We used R version 4.1.1 [51] for all R packages. We applied the Bioconductor package edgeR [52] originally developed for RNA-seq data and later adapted for methylation data of mammalian genomes, on the more complex plant methylome. In mammals, the vast majority of methylated cytosines occur in CpG context, while for plants methylation occurs in all three sequence contexts (CpG, CHG, CHH). Identified methylated sites were first assigned to relevant predefined regions with the R-package GenomicRanges and methylated regions were identified with the package edgeR v3.40.2. We focus on gene bodies and promoters as predefined regions to limit the number of pairwise comparisons in the next steps. We defined gene-body methylation (gbM) as enrichment of CpG-only methylation within gene bodies with a depletion of methylation at transcription start site (TSS) and transcription termination sites (TTS) [6]. Promoters were defined as the region between 2000 bp upstream and 200 bp downstream the TSS. Unlike most other approaches to methylation sequencing data, the edgeR workflow keeps the counts for methylated and unmethylated reads as separate observations. Generalized linear models are used to fit the total read count (methylated plus unmethylated) at each genomic locus, in such a way that the proportion of methylated reads at each locus is modeled indirectly as an over-dispersed binomial-like distribution. Keeping methylated and unmethylated read counts as separate data observations allows the variability of the data to be modeled more directly and perhaps more realistically [52]. We identified DMRs by grouping the WGBS data from the DNA samples of individual plants by their corresponding parent-of-origin (ortet located in Hungary, Italy, Spain and the UK, respectively). This resulted in four groups with three (Hungary), four (Italy), five (Spain) and four (UK) biological replicates per group. We identified statistically significant DMRs in between-group pairwise comparisons for two pre-defined regions (genes and promoters) for each of the three sequence contexts (CpG-, CHG, CHH-DMRs), resulting in 36 pairwise comparisons. Additional file 9 includes the R-script with the code to reproduce this analyses.

#### 5.1.5 Identification of *de novo* differentially methylated regions

In addition to identifying DMRs in promotors and gene bodies, we also scanned the methylomes to detect unbiased *de novo* DMRs. This was done using the Bioconductor package dmrseq v1.18.0 [23]. A DMR was called differentially methylated when the adjusted p-value was smaller or equal than 0.05. Lastly, the position of the differentially methylated DMRs was compared to the current genomic annotation. We also applied this package developed for mammalian epigenomes to the more complex plant epigenome. In most former plant studies, computational approaches for detecting DMRs do not provide accurate statistical inference. Methods to identify DMRs in WGBS experiments are greatly hindered by the high-dimensionality and low sample size setting that is common in high-throughput genomics studies. The number of tests performed is equal to the number of loci analyzed, which is very large in typical WGBS studies. The approach implemented in dmrseq improves the specificity and sensitivity of lists of regions and accurately controls the false discovery rate (FDR) [23]. Dmrseq uses a two-stage approach. First, candidate regions are detected and afterwards their statistical significance is evaluated. In brief, dmrseq starts by smoothing the signal, because neighboring cytosines are highly correlated. Subsequently, candidate regions are defined by segmenting the genome groups of cytosines that show consistent evidence of differential methylation. In the second stage, a statistic is computed for each candidate region. This statistic takes into account variability between biological replicates and spatial correlation among neighboring loci [23]. To our knowledge, this is one of the first studies applying this approach on plant genomes. Only cytosines that have coverage in all samples were retained.

To investigate whether the epigenetic variation within vegetative offspring in a non-parental environment correlates with the parental climate, we used the i) WGBS data of the 13 pooled DNA samples and ii) the WGBS data of the individual ramets. For the pooled samples, we grouped the WGBS data in three groups based on the average temperature in January (JAN) at the geographic location of the Lombardy poplar parent-of-origin. The geographic location of the ortets included in the analysis of the pooled samples is given in Figure 1B. We used JAN as the representative climate variable. Temperature is an important environmental factor determining plant vitality and growth [53]. Cold winters effect tree dormancy and soil water availability and January is generally the coldest month of the year at locations of the sampled Lombardy poplars. The climate variable JAN is correlated with the average annual precipitation (correlation coefficient: 0.6; see Additional file 7), which is another important climate variable determining plant growth. We used data from WorldClim 2 [54] for the period 1965 – 2015 to characterize the home environment of each ortet. In addition to the 13 pooled samples, we included one individual sample (WGBS04, sample from Spain, Tree_ID SPC1) in this analysis. This resulted in three groups with five (group 1, range JAN (°C); 1.82 - 3.61), six (group 2, range JAN (°C); −1.40 - 0.15) and three (group 3, range JAN (°C); 4.97 - 8.0) biological replicates, respectively (Figure 1B). We searched for statistically significant *de novo* DMRs in between-group pairwise comparisons for each of the three sequence contexts (CpG-, CHG, CHH-DMRs), resulting in nine pairwise comparisons. We performed a similar analysis for the individual ramets sampled at the begin (March) and in the middle (June/July) of the growing season, by grouping the WGBS data based on the average January temperature at the geographic location of the Lombardy poplar parent-of-origin (i.e. ramets originated from the ortets from Hungary and UK versus ramets from the ortets located in Spain and Italy).

#### 5.1.6 Gene ontology enrichment analysis

To test which biological processes were over-represented in sequences containing DMRs (as compared to the *Populus trichocarpa* genome), we performed a statistical overrepresentation test in PANTHER Classification System version 16 (http://pantherdb.org/) [55]. We performed a Fisher’s exact test with FDR correction using the differentially methylated gene features (genes IDs and promoters) as input data and the *Populus trichocarpa* GO-Slim annotation data as reference data, with a p-value cutoff of 0.05. We further explored the biological importance of the list of genes in significant DMRs with the Bioconductor package simplifyEnrichment v1.8.0 [56]. This package implements the binary cut algorithm for clustering similarity matrices of functional GO terms. GO terms that were significant over-represented in genes containing DMRs are clustered and their similarities are visualized in a heatmap. The package simplifyEnrichment visualizes the summaries of clusters by word clouds which helps to interpret the common biological functions shared in the clusters. Additional file 10 includes the R-script with the code to reproduce the clustering and visualizing of the GO enrichment results.

#### 5.1.7 Cluster analysis

We performed a clustering analysis based on the methylated cytosines from DNA samples of individual plants to investigate the methylome variability between plants and the stability of the epigenome during a growing season and across vegetatively reproduced generations. We first explored the overall differences in cytosine methylation levels (differential methylated positions, DMPs) using a hierarchical clustering analysis based on the Pearson’s correlation distance and the Ward method (minimum variance method) with the Bioconductor package methylKit 1.24.0. [57]. We used the filtering function ‘filterByCoverage’ as described in the vignette of methylKit, i.e. removing loci with lower than 10x coverage and loci in 99.9th percentile of coverage in each sample. We further investigate DMPs between the samples in promotors and gene bodies using multi-dimensional scaling (MDS) plots based on the M-value for each of the three sequence contexts (CpG, CHG, CHH) separately with the R package edgeR. The M-value is a common measure of the methylation level and is defined as M = log2((Me + α)/(Un + α)) where Me and Un are the methylated and unmethylated intensities and α is some suitable offset to avoid taking logarithms of zero. In the MDS plot the distance between each pair of samples represents the average logit change between the samples for the top most differentially methylated loci between that pair of samples.

### 5.2 Common environment experiment

#### 5.2.1 Experimental design

The design of the common environment experiment is described in Vanden Broeck et al. (2018). In brief, during the winter of 2016 - 2017 dormant one-year old twigs were collected on 94 adult putative Lombardy poplar trees (hereafter called; ortets, F_0_-generation) located at 37 different sites in Europe and Asia, including those that were used in the WGBS analysis. The twigs were shipped by express mail to the Research Institute for Nature and Forest (INBO) in Geraardsbergen, Belgium and stored in the fridge at 4°C until the establishment of the greenhouse experiment in March 2017. Out of the 94 putative Lombardy poplar trees, 72 were identified as “true” Lombardy poplars based on the multilocus genotypes obtained with 11 nuclear microsatellite polymorphisms. The poplar trees identified as “true” Lombardy poplars include the most common genotype that was shared by most of the sampled trees (G01; see Vanden Broeck et al. 2018) and two other multilocus genotypes (G07 and G14) that differed from the most common genotype for only one out of the 22 alleles at locus WPMS05 (mismatch of one repeat).

In March 2017 up to 14 (mean: 12.8, range: 4 – 14) cuttings (hereafter called; ramets, F_1_-generation) per donor tree were planted in trays filled with standard potting soil and were grown under similar light and temperature conditions in an open greenhouse at INBO. After the first growing season, the ramets were cut back and new shoots resprouted on two-year old roots in spring 2018. After the second growing season, in February 2019, the ramets were transplanted from the trays to larger, single pots (volume ∼ 10L) with potting soil (50% white peat / 50% black peat, 0.12% nitrogen, 0.14% phosphorous, 0.24% potassium) and the experiment was transferred 30 km north to an open container field located in Melle, Belgium (lat. 50,7000556°, lon. 3.792222°). Also here, the plants were grown under similar temperature and light conditions for two more growing seasons (2019 and 2020) and were regularly watered.

#### 5.2.2 Bud set observations

We combined the previously published bud set observations of 2017 with new observations of three consecutive growing seasons, resulting in a dataset spanning four growing seasons (2017-2020). In total, 793 ramets from 67 adult Lombardy poplar ortets from 28 locations and 13 countries were included in the experiment. These ortets were located across a 2117 km-wide latitudinal transect from Dalkeith (Scotland) to Salermo (Italy) and across a 1275 km-wide longitudinal gradient from Loiret (France) to Nagyvenyim (Hungary). In poplar, latitude is related to the length of the growing season as demonstrated by differences in growth cessation in autumn [48]. An interactive map of the location of the ortets is available at [Additional file 7](https://zenodo.org/deposit/7400979).

Bud set was scored from summer to autumn, on the apical bud of a 1-yr-old shoot on 1-yr-old (2017), 2-yr-old (2018), 3-yr-old (2019) and 4-yr-old (2020) roots. We used a seven stage scoring system to cover onset and duration of bud set developed for *Populus nigra* by Rohde et al. [48]. Scores go from 3 (growing apical meristem) to 0 (fully developed bud), in 0.5 intervals. Observations were performed once a week starting in July (except for 2017 where observations started at the beginning of August) until all plants had fully developed apical buds. Differences in the ontological status of the ramets, the original nutrient condition of the ramets and/or micro-environmental variation may be associated with the physiological plant state which may affect the process of bud set formation. Plant growth is a good indicator for the physiological plant state in pot experiments [58]. We measured the length of the last year’s shoot (± 1cm) in January-February 2019 and February 2020. During each growth period, the physiological condition of the plants in the common environment experiment was visually monitored in order to define the need for watering or pest control. Fertilization, water availability and plant pests were controlled. Temperature modifies the sensitivity to day-length signals at growth cessation and influences the duration of bud formation in poplar [48]. Data on the daily minimum temperature during plant growth was obtained from a nearby weather station (Hoeilaart, lat. 50,77673°, lon. 4.46833°). After inspection of the plants’ physiological state for statistical outliers for the response variable (bud set scores), none of the outliers were removed from the data. In total and over the 4-yr period, we observed bud set on 785 ramets (on average, range 791 −784) during 46 different dates, with a mean of 11.5 (range; 8 - 16) observation dates per year, resulting in 35,950 data points.

#### 5.2.3 Data analysis

Using R version 4.1.1 (R Core Team, 2021) and the package glmmTMB v1.1.5 [59] we performed a linear mixed effects analysis of the relationship between bud set phenology of the ramets in the common environment and the latitude of origin of the home site of the Lombardy poplar parent-of-origin. Bud set observations over the different years were considered as independent. To account for non-independence of the bud set scores of the ramets from a single ortet, we aggregated the data for the ramets per ortet by averaging the bud set scores per ortet. The different ortets (parents-of-origin) are represented by the variable ID_TREE. We used the variables ID_TREE nested within Location as random intercepts for the model. This accounts for the correlation between ortets (ID_TREE) of a single location. To account for correlation in the time series of bud set scores (they can be considered as stationary series), we use an autoregressive function (AR(1)). An AR(1) model estimates the correlation between two observations in adjacent time steps. The following fixed effects (with interaction terms) were included into the selected model; the temperature sum of daily minimum temperature at the common environment experiment calculated from 1 July (Tsmin), the latitude of origin of the parent-of-origins (Lat, as a proxy for climate and photoperiod) and the year of observation (Year). The year of observation (Year) was included as a factor variable (four levels), the other predictor variables were continuous variables. The year 2017 was set as the reference year. The variable Tsmin was centered and standardized (called Tsminc), and the variable Lat was centered on 50 (range −9.2 to 5.8, unit = 1 degree). For the sake of simplicity, only two-way interactions were taken into account. The significance (p-value) of fixed effects was tested by likelihood ratio tests of the full model with the effect in question against the model without the effect in question (null-model) using ANOVA. To avoid overfitting and for simplicity, we also used the Bayesian information criterion (BIC) in addition to the likelihood ratio tests, to select the optimal model.

The model with the best fit was, in R syntax, as follows;

Score ∼ Tsminc + I(Tsminc^2) + Lat + fYear + (1 | Location/ID_TREE) + (ar1(times + 0 | fYear)) + Tsminc:fYear + I(Tsminc^2):fYear + Lat:fYear

## 6 Declarations

### 6.1 Ethics approval and consent to participate

Not applicable.

### 6.2 Consent for publication

Not applicable.

### 6.3 Availability of data and materials

The datasets supporting the conclusions of this article are available in the Zenodo repository, https://doi.org/10.5281/zenodo.7400979 [60].

### 6.6 Author Contributions

AVB, KC, BH and FVN designed the experiments. EDM performed the WGBS library preparation. TM performed the WGBS data analysis, PV performed the bud set data analysis. AVB, TM and PV wrote the manuscript and prepared the figures. KC, BH, DD and FVN revised the manuscript and provided input. All authors read and approved the final manuscript. AVB and TM contributed equally to this work.

## 6.7 Acknowledgements

The authors are very thankful to Afrooz Alimohamadi, Farhad Asadi, Mohsen Calagari, Jan Douda, Arion Turcsán, István Bach, Bert Maes, Joukje Buiteveld, Cornelis Van Oosten, Dalibor Ballian, Davorin Kajba, Eduardo Notivol Paino, François Lefèvre, Marc Villar, Georg von Wühlisch, Gregor Božič, Ilka Yonovska, Lorenzo Vietto, Nicolae Talagai, David Halfmaerten and Stephanos Diamandis for providing plant material and background information. The authors also thank Nico De Regge, Marc Schouppe, Wim De Clercq and Kurt Schamp for their help in the bud set observations, and Serge Goossens and Stefaan Moreels for taking care of the plants in the greenhouse experiment. We also thank Sabrina Neyrinck for the laboratory analyses.

## 11 Additional files

- **Additional file 1.** CSV-file with information on the Lombardy poplar trees samples used for whole genome bisulfite sequencing (WGBS) in the two methylome experiments.
- **Additional file 2.** Zip-folder with HTML-files of the mapping and methylation statistics of the genomes of the 16 individual Lombardy poplar samples for the three sequence contexts (CpG, CHG, CHH) (bismark reports).
- **Additional file 3**. Zip-folder with: i) Excel-files listing the genes in DMRs, and ii) PNG-files with the ‘Biological Coefficient of Variation (BCV)’-plots between any of the six pairwise comparisons of Lombardy poplars grouped per ortet and identified with Bioconductor package edgeR. DMRs were identified between groups by grouping the WGBS data from 16 individual Lombardy poplar ramets by their corresponding parent-of-origin (‘HUN4’ located in Hungary, ‘ITS3’ in Italy, ‘SPC1’ in Spain and ‘UKD2’ in the UK).
- **Additional file 4**. Excel-file with the total list of GO terms that were enriched in DMRs. DMRs were identified between groups by grouping the WGBS data from 16 individual Lombardy poplar ramets by their corresponding parent-of-origin (ortet ‘HUN4’ located in Hungary, ‘ITS3’ in Italy, ‘SPC1’ in Spain and ‘UKD2’ in the UK).
- **Additional file 5**. Zip-folder with PNG-files representing heatmaps and Excel spreadsheets with clustered GO terms significant over-represented in promoters and gene regions located in DMRs. DMRs were identified between groups by grouping the WGBS data from 16 individual Lombardy poplar ramets by their corresponding parent-of-origin (ortet ‘HUN4’ located in Hungary, ‘ITS3’ in Italy, ‘SPC1’ in Spain and ‘UKD2’ in the UK. The files were obtained with the Bioconductor package simplifyEnrichment.
- **Additional file 6.** CSV-file with the raw data of the bud set observations in the common garden experiment.
- **Additional file 7.** HTML-file with the R source codes to reproduce the results of the bud set analysis.
- **Additional file 8.** A text-file representing the Snakefile (i.e. a readable Python-based workflow) including the different steps and rules of the bioinformatics of the WGBS data analyses.
- **Additional file 9.** RMD-file with the code to reproduce the analyses to identify differential methylated predefined regions.
- **Additional file 10.** R-script with the code to reproduce the clustering and visualizing of the GO enrichment results.

